# 28S-rsRNA facilitates PASMC proliferation through PKM2 β-hydroxybutyrylation

**DOI:** 10.64898/2026.11.09.765510

**Authors:** Aijing Liu, Yingqi Wang, Yu Pan, Jimiao Zhang, Chunmiao Liu, Xueyin Liao, June Bai, Hanliang Sun, Xiaoxu Zhao, Zhitu Bao, He Zhu, Yunyi Ji, Yiming Xu, Yan Xing, Haiyang Tang, Daling Zhu, Hang Yu

## Abstract

Ribosomal RNA-derived small RNAs (rsRNAs) are nucleic acid fragments originating from rRNA, the functions and roles of which in diseases remain incompletely understood. We identified 28S rsRNA-CTC_L20, an rsRNA whose expression is upregulated in hypoxic pulmonary artery smooth muscle cells (PASMCs) and in lung tissues of pulmonary hypertension (PH) models. In hypoxic cells, 28S rsRNA-CTC_L20 promotes PASMC proliferation, and its knockdown mitigates SuHx-induced pulmonary hypertension and vascular remodeling. Mechanistically, 28S rsRNA-CTC_L20 modulates β-hydroxybutyrate (BHB) production by maintaining the stability of ketogenic enzyme mRNAs through its “CTGNNGG” motif. Elevated levels of BHB enhance the β-hydroxybutyrylation modification at K166 of pyruvate kinase M2 (PKM2), increasing its activity and driving glycolysis to sustain proliferation. These results link ketone body metabolism to glycolysis in hypoxic cells and define an rsRNA-BHB-PKM2 β-hydroxybutyrylation axis in PH. They also point to 28S rsRNA-CTC_L20 as a potential target for treating vascular remodeling. **Keywords:** 28S rsRNA-CTC_L20; β-hydroxybutyrylation modification; Hypoxia; pulmonary artery smooth muscle cells; pulmonary hypertension

## Introduction

Pulmonary hypertension (PH) is a heterogeneous group of cardiopulmonary disorders characterized by a particularly poor prognosis, despite current vasodilator therapies^1^. In PH, pulmonary artery smooth muscle cells (PASMCs) switch from a quiescent contractile phenotype to a synthetic, proliferative one, and this switch drives medial hypertrophy and distal arteriolar occlusion^2^. Hypoxia, the strongest environmental trigger of PH, amplifies PASMC proliferation, and noncoding RNA networks contribute to this response^3^.

A Warburg-like aerobic glycolysis, which supplies ATP and biosynthetic precursors for PASMC expansion, is a consistent feature of the PH metabolic phenotype. Platelet-derived TGF-β1 enhances the aerobic glycolysis of PASMCs through PKM2 (pyruvate kinase M2) upregulation, and genetic or pharmacologic interference with this axis attenuates SuHx-induced PH^4^. Consistent with this, loss of circSMOC1 promotes metabolic reprogramming and PASMC glycolysis via the PTBP1/PKM2 axis^5^, while direct PKM2 activation blunts hypoxia-induced PASMC proliferation and migration^6^. PKM2-dependent glycolysis is therefore a potential target in vascular remodeling.

PKM2 activity is governed by multiple post-translational modifications (PTMs). One of these, lysine β-hydroxybutyrylation (Kbhb), is a metabolite-derived acylation driven by the ketone body β-hydroxybutyrate (BHB), and its role in PH has not been examined. Kbhb was described only recently, and its regulatory enzymes (p300 as writer; HDAC1/2 as erasers) and thousands of substrate sites have been mapped in mammalian cells^7^. Proteomic mapping further revealed that BHB-induced Kbhb is enriched on enzymes of glycolysis/gluconeogenesis and pyruvate metabolism, directly linking this modification to glycolytic control^8^. PKM2 is itself subject to many PTMs that tune its dimer–tetramer equilibrium and nuclear functions^9^, but whether Kbhb of PKM2 licenses glycolytic reprogramming in hypoxia is unknown.

rsRNAs (rRNA-derived small RNAs) are a distinct class of regulatory noncoding RNAs produced by precise cleavage of ribosomal RNAs, yet their functions in cardiovascular disease are largely unknown. Early profiling showed that 5′/3′ rRNA fragments (rRFs), including 28S-derived species, are abundant, length-graded small RNAs rather than degradation by-products, and that they modulate proliferation and apoptosis^10^, and a defined sequence of 28S rRNA-derived small RNAs is broadly enriched across human somatic tissues and associates with inflammation^11^. An rsRNA-28S species binds target mRNAs and regulates cancer cell phenotypes^12^, so 28S rsRNAs can act as sequence-specific post-transcriptional regulators. Whether hypoxia induces 28S rsRNAs that govern pulmonary vascular cell metabolism has not been investigated.

Ketone body metabolism provides an additional link to glycolysis. BHB is no longer viewed only as an alternative fuel; it also acts as a signaling metabolite that relaxes pulmonary arteries and improves hemodynamics in patients with PH^13^. In PAH, fatty acid oxidation generates BHB, which sensitizes TRPV4 channels in lung microvascular endothelial cells and links metabolism directly to function in the pulmonary circulation^14^. Systemically, ketone bodies have broad vasculoprotective and signaling effects, including protein β-hydroxybutyrylation, that are now being developed as cardiovascular therapeutic targets^15^. How ketone body production is regulated in hypoxic PASMCs, and whether it communicates with glycolysis, remains unknown.

Several hypoxia-inducible noncoding RNAs shape PASMC fate: loss of lncRNA Rps4l exacerbates hypoxic PASMC proliferation via ILF3/HIF-1α^3^; circPMS1 is induced under hypoxia and drives proliferation of PASMCs, endothelial cells and pericytes^16^; lncRNA PVT1 upregulation promotes autophagy deregulation and PASMC growth^17^. We screened hypoxia-responsive small RNAs and identified 28S rsRNA-CTC_L20, a 28S rRNA-derived fragment whose expression is selectively elevated in hypoxic PASMCs and in lung tissues from PH models. On this basis, we hypothesized that a hypoxia-induced rsRNA orchestrates ketone body metabolism to enable glycolytic reprogramming and proliferation in PASMCs.

We show here that 28S rsRNA-CTC_L20 is upregulated under hypoxia and promotes PASMC proliferation, whereas its deficiency attenuates SuHx-induced PH and vascular remodeling. Mechanistically, 28S rsRNA-CTC_L20 stabilizes ketogenic enzyme mRNAs through its “CTGNNGG” motif, raising BHB levels that enhance Kbhb at PKM2-K166, increase PKM2 activity and activate glycolysis to sustain proliferation. This work connects ketone body metabolism to glycolysis in hypoxic cells and defines an rsRNA–BHB–PKM2(K166^bhb^) axis in PH. By identifying 28S rsRNA-CTC_L20 as a regulator of vascular remodeling, our findings point to a potential therapeutic target for PH.

## METHODS and MATERIALS

### Data Availability

The authors declare that all supporting data are available within the article and its Supplemental Material. Additional deidentified data, analytic methods, and study materials are available from the corresponding author upon reasonable request. Please see the Major Resources Table in the Supplemental Materials.

### Study Approval

Animal protocols were approved by the Medical Animal Ethics Committee of Harbin Medical University–Daqing Campus (approval No. HMUDQ20260122001) and conformed to the NIH Guide for the Care and Use of Laboratory Animals (NIH Publication No. 85-23, revised 1996); the study is reported in accordance with the ARRIVE guidelines. Human lung tissue and plasma were obtained after written informed consent, with approval of the Ethics Committees of The First Affiliated Hospital of Guangzhou Medical University and Harbin Medical University–Daqing Campus, in accordance with the Declaration of Helsinki. Hang Yu had full access to all the data and takes responsibility for their integrity and the data analysis.

### Human Samples

Lung tissue was obtained from 3 patients with hypoxia-related PH (secondary to COPD) undergoing lung transplantation, with the diagnosis confirmed by right heart catheterization and no prior PH-targeted therapy, and from 3 unused donor lungs. Plasma was collected from patients with PH and from healthy volunteers between 2022 and 2024; blood was centrifuged at 1500 g for 15 min at 4°C and plasma was stored at −80°C. Clinical characteristics are given in Tables S1 and S2 (see supplemental matetials). All subjects were Asian; because of the small sample size, sex- and ethnicity-specific analyses were not performed and the human data are regarded as preliminary.

### Experimental Animals

Male C57BL/6J mice (6∼8 weeks, 18∼22 g; Experimental Animal Department of Harbin Medical University) were acclimated for 7 days with free access to food and water under a 12:12-hour light–dark cycle. Only male mice were used because estrogen modifies the severity of hypoxia-induced pulmonary hypertension; this limitation is acknowledged.

### SuHx Model and Interventions

Mice were randomly allocated to normoxic control or SuHx groups. SuHx PH was induced by normobaric hypoxia (FiO_2_ 0.10) for 21 days with intraperitoneal Sugen5416 (20 mg/kg) on days 1, 7, and 14, followed by return to normoxia (FiO_2_ 0.21). Two weeks later, mice received a single intranasal instillation of AAV6 carrying a 28S-rsRNA-CTC_L20 antisense oligonucleotide (AAV6-ASO) or a scrambled control (1×10^10^ to 1×10^11^ vector genomes in 20∼30 µL Hanks’ Balanced Salt Solution; GeneChem, Shanghai, China) and were studied 3 weeks thereafter. The bosentan group was included and received bosentan (10 mg/kg/day) by oral gavage for 3 weeks under normoxia (21% O_2_). All invasive procedures were performed under tribromoethanol anesthesia.

### Echocardiography and Hemodynamics

Under anesthesia (tribromoethanol, 60 mg/kg intraperitoneal), pulmonary artery acceleration time (PAAT) and pulmonary artery velocity-time integral (PAVTI) were measured with a Vevo 2100 system. Right ventricular systolic pressure (RVSP) was then recorded with a 1.2 French catheter (Scisense, Inc.) and an FA-404 recorder, and right ventricular hypertrophy was expressed as RV/(LV+septum). Operators were blinded to group allocation.

### Cell Culture and Hypoxia

Mouse pulmonary artery smooth muscle cells (mPASMCs) and pulmonary artery endothelial cells (mPAECs) (Hunan Fenghui Biotechnology, Changsha, China) were cultured in smooth muscle or endothelial cell medium at 37 °C in 5% CO_2_, used between passages 3 and 8, and were mycoplasma-negative. Hypoxia was produced in a tri-gas incubator (Heal Force) with 3% O_2_, 5% CO_2_, and 92% N_2_; normoxic controls were kept at 21% O_2_.

### Small RNA Profiling

Because conventional small RNA sequencing underestimates sncRNAs bearing modifications that block adapter ligation and reverse transcription, small RNAs from hypoxic and normoxic mPASMCs (n=3 per group) were profiled by PANDORA-seq, which removes these modifications before library construction. Reads were annotated against rRNA, tRNA, and other noncoding RNA references; differentially expressed sncRNAs were defined as |log2 fold change| ≥ 1 with adjusted *P < 0.05*. Putative targets of 28S-rsRNA-CTC_L20 were predicted from sequence complementarity, and the predicted target genes were analyzed for Gene Ontology and KEGG enrichment with Metascape.

### RNA Quantification

Total RNA was extracted with TRIzol, cytoplasmic and nuclear RNA were separated with a PARIS kit. After reverse transcription (PrimeScript RT reagent Kit), quantitative PCR was run on a LightCycler 480 II system with TB Green Premix Ex Taq; β-actin, 18S, and U6 served as references for total, cytoplasmic, and nuclear RNA, and relative expression was calculated by the 2^−ΔΔCt^ method.

### Protein Analyses

Proteins were separated on 10%∼12% polyacrylamide gels, transferred to nitrocellulose, and detected with primary antibodies, horseradish peroxidase-conjugated secondary antibodies, and enhanced chemiluminescence. For co-immunoprecipitation, lysates were pre-cleared with Protein A/G magnetic beads and incubated with a PKM2 antibody. β-hydroxybutyrylation (Kbhb) of PKM2 was assessed by immunoprecipitating PKM2 and immunoblotting with a pan-Kbhb antibody, and the modified residue was identified by liquid chromatography–tandem mass spectrometry. Antibody details are provided in the Major Resources Table. Each band shown derives from an independent biological sample, and no bands were spliced or otherwise manipulated.

### Proliferation, Cell Cycle, and Viability

Cell viability was measured with a Cell Counting Kit-8 assay, DNA synthesis by EdU incorporation (10 µM, 2 h) with Click detection, proliferation by Ki67 immunofluorescence, and cell cycle distribution by propidium iodide staining and flow cytometry.

### Ketone Body Measurements, mRNA Stability, and Histology

Intracellular β-hydroxybutyrate and acetoacetate were measured in mPASMC lysates with commercial assays. For mRNA stability, cells at 70%∼80% confluence were exposed to actinomycin D (2 µg/mL), harvested at serial time points, and the decay rate and half-life were derived by fitting a one-phase decay model. Lung sections (5 µm) were stained with hematoxylin–eosin and Masson trichrome and quantified by observers blinded to group allocation; 28S-rsRNA-CTC_L20 was localized by fluorescence in situ hybridization with a biotin-labeled probe and SA-Cy3.

### Randomization and Blinding

Mice were allocated to groups in a randomized manner. Investigators performing echocardiography, hemodynamics, and histological quantification, and those analyzing images, were blinded to allocation; cell culture samples were processed and analyzed in random order.

## Statistical Analysis

Data are expressed as mean±SD of varying numbers of independent biological replicates, where n denotes independent animals or independent cell preparations rather than technical replicates. Normality was assessed with the Shapiro–Wilk test. Two groups were compared with an unpaired 2-sided Student t test, with Welch’s correction when variances were unequal; 3 or more groups were compared with 1-way analysis of variance followed by Tukey post hoc test. Non-normally distributed data were analyzed with the Mann–Whitney U or Kruskal–Wallis test. Analyses were performed with GraphPad Prism (version 10.2.3). Tests were 2-sided and exact P values are reported; *P<0.05* was considered statistically significant, and P values were adjusted for the multiple comparisons within the Tukey post hoc test where applicable; All experiments were independently repeated at least 3 times.

## Results

### 28S-rsRNA-CTC_L20 is induced by hypoxia in PASMCs and enriched in the pulmonary arteries of SuHx mice and patients with PH

To identify small non-coding RNAs (sncRNAs) involved in pulmonary hypertension, we performed PANDORA-seq. Among the differentially expressed sncRNAs, 28S-rsRNA-CTC_L20 was one of the most significantly upregulated candidates (Fig. 1A). Sanger sequencing verified the sequence of 28S-rsRNA-CTC_L20, and agarose gel electrophoresis confirmed the size and uniqueness of the amplified product (Fig. 1B). qRT-PCR analysis showed that the expression of 28S-rsRNA-CTC_L20 was significantly increased in the lung tissues of SuHx mice, whereas no significant change was detected in the heart, liver, spleen or kidney (Fig. 1C). Consistently, the level of 28S-rsRNA-CTC_L20 was elevated in the blood of SuHx mice (Fig. S1A). FISH analysis further showed that 28S-rsRNA-CTC_L20 was upregulated in the α-SMA-positive pulmonary arteries of SuHx mouse lung tissues (Fig. 1C). We next determined the cellular source of 28S-rsRNA-CTC_L20. In mPASMCs exposed to hypoxia for 0, 24, 48 and 72 h, the expression of 28S-rsRNA-CTC_L20 increased with the duration of hypoxia (Fig. S1B). Hypoxia increased the expression of 28S-rsRNA-CTC_L20 in mPASMCs, whereas no significant change was observed in mPAECs under the same conditions (Fig. 1D). qRT-PCR analysis of the nuclear and cytoplasmic fractions, together with FISH analysis, showed that 28S-rsRNA-CTC_L20 was distributed in both the nucleus and the cytoplasm of mPASMCs, with U6 and 18S rRNA as the nuclear and cytoplasmic controls, respectively, and that it shifted toward the cytoplasm under hypoxia (Fig. 1E, S1C). The expression of 28S-rsRNA-CTC_L20 was also elevated in the plasma of patients with PH compared with healthy volunteers, and FISH analysis showed its upregulation in the α-SMA-positive pulmonary arteries of lung tissues from patients with PH (Fig. 1F).

**Figure 1.**
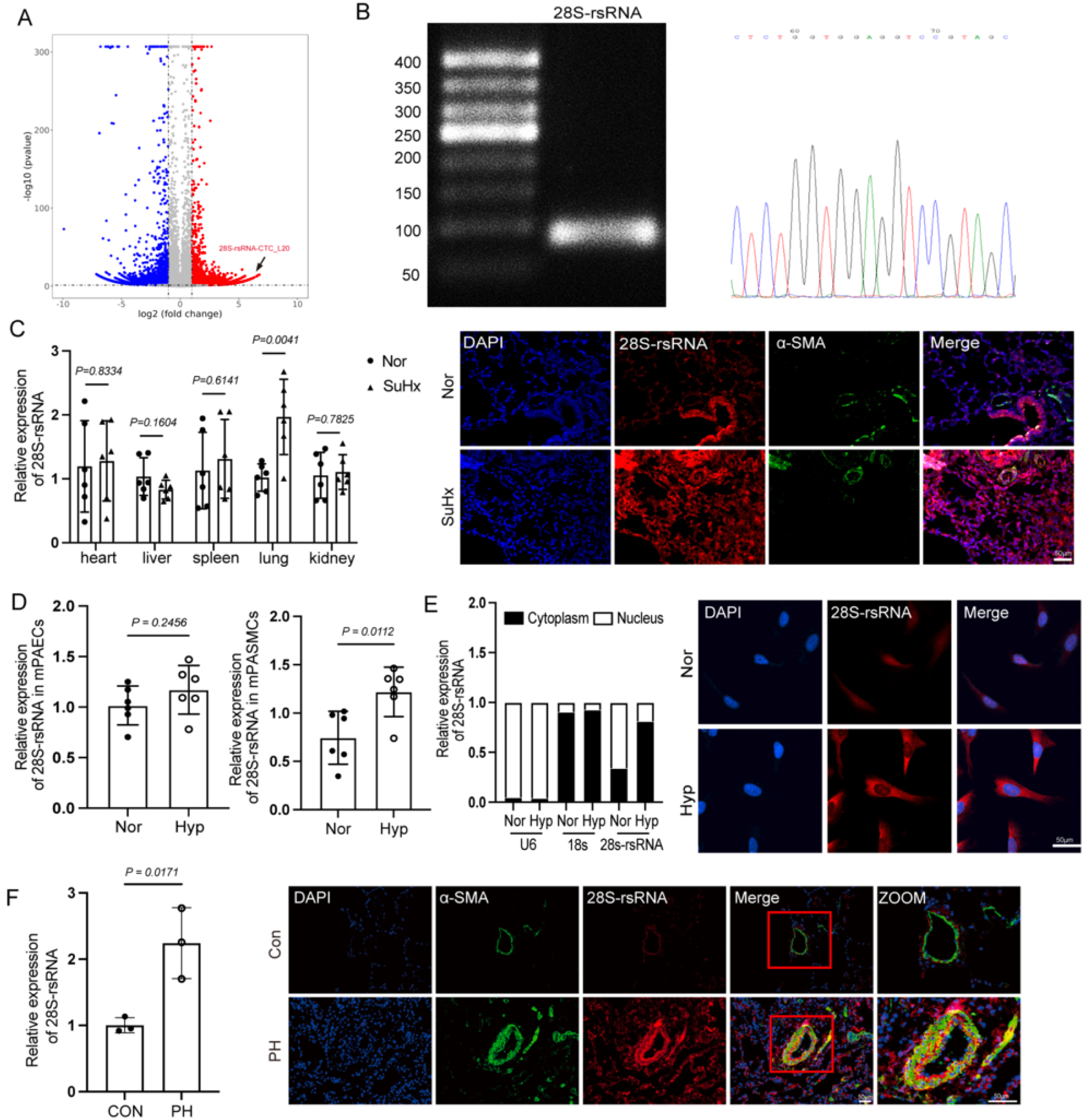
28S-rsRNA-CTC_L20 is induced by hypoxia in PASMCs and enriched in the pulmonary arteries of SuHx mice and patients with PH. (A) Volcano plot of PANDORA-seq data on small noncoding RNAs from mPASMCs; upregulated and downregulated transcripts appear in red and blue, and the arrow points to 28S-rsRNA-CTC_L20. (B) The amplified 28S-rsRNA-CTC_L20 product was verified by Sanger sequencing and resolved on an agarose gel. (C) qRT-PCR in heart, liver, spleen, lung and kidney of normoxic (Nor) and SuHx mice (n=6 mice per group), and FISH of lung sections for 28S-rsRNA-CTC_L20 (red) and α-SMA (green) with DAPI counterstain (n=3 mice per group). Scale bar, 50 μm. (D) mPAECs and mPASMCs cultured under normoxia (Nor) or hypoxia (Hyp) (n=6 independent cultures per group). (E) Nuclear and cytoplasmic fractions of mPASMCs, with U6 and 18S rRNA as compartment markers (n=3 independent experiments); FISH images are shown alongside. Scale bar, 50 μm. (F) Plasma from healthy volunteers (CON) and patients with PH (n=3 subjects per group), and FISH of α-SMA-positive pulmonary arteries in human lung. Scale bar, 50 μm. Data are mean±SEM. α-SMA indicates alpha-smooth muscle actin; FISH, fluorescence in situ hybridization; mPAEC, mouse pulmonary artery endothelial cell; mPASMC, mouse pulmonary artery smooth muscle cell; PANDORA-seq, panoramic RNA display by overcoming RNA modification aborted sequencing; PH, pulmonary hypertension; SuHx, SU5416 plus hypoxia.

### Silencing 28S-rsRNA-CTC_L20 blunts hypoxia-driven proliferation of mPASMCs and its overexpression is sufficient under normoxia

To explore the function of 28S-rsRNA-CTC_L20, we performed GO biological process (GO:BP) enrichment analysis of its predicted target genes. The enriched terms included RNA splicing and proliferation-related processes, among them the regulation of smooth muscle cell proliferation (Fig. 2A). Cell cycle analysis by flow cytometry showed that hypoxia promoted the progression of mPASMCs into S phase, and this effect was inhibited by the 28S-rsRNA-CTC_L20 inhibitor (Fig. 2B, S2A). In hypoxic mPASMCs, the 28S-rsRNA-CTC_L20 inhibitor decreased the expression of PCNA, CDK1, CDK2, Cyclin A and Cyclin E (Fig. 2C, S2B), reduced the proportion of EdU- and Ki67-positive cells (Fig. 2D) and reduced cell viability (Fig. 2E).

**Figure 2.**
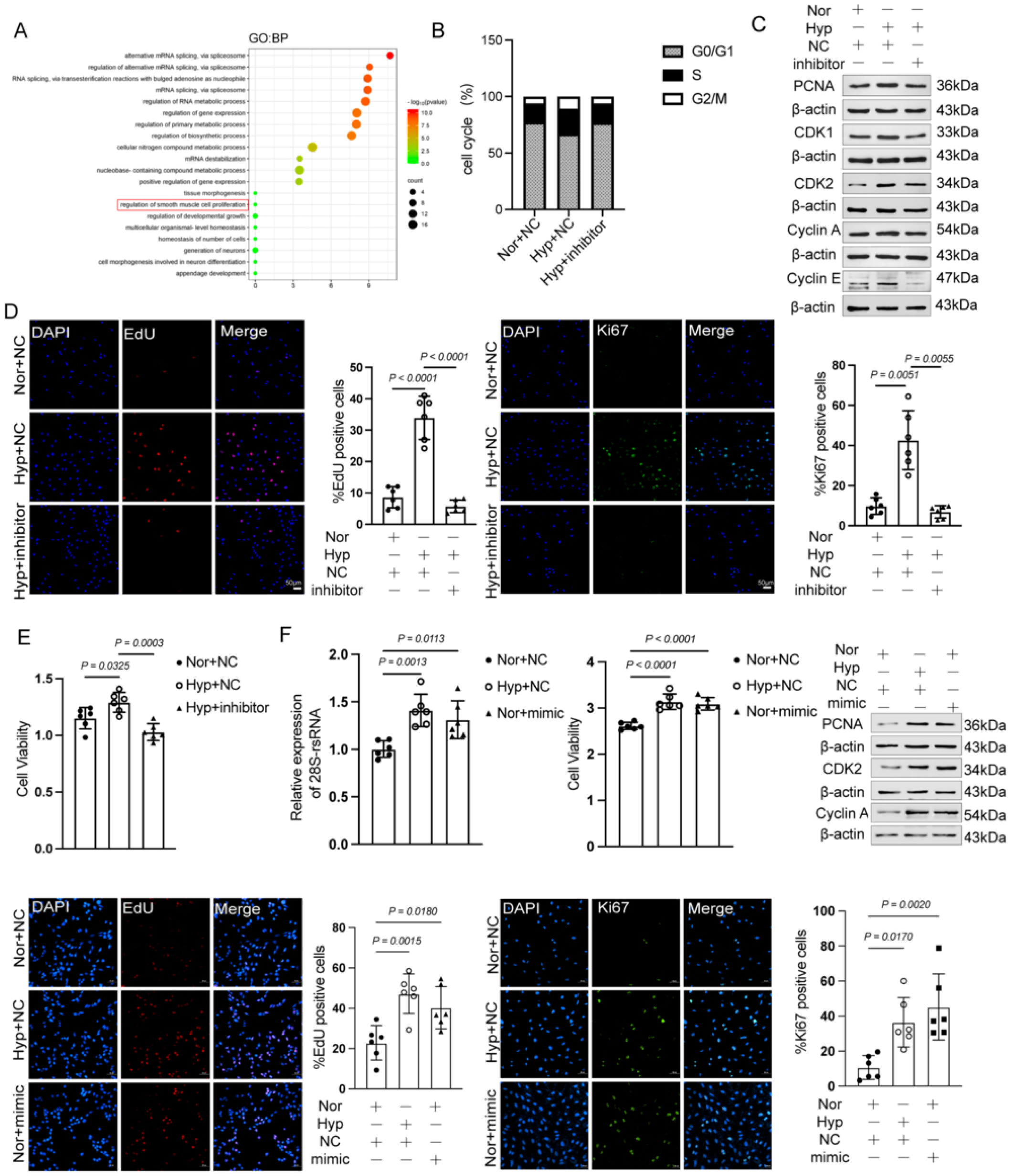
Silencing 28S-rsRNA-CTC_L20 blunts hypoxia-driven proliferation of mPASMCs, and its overexpression is sufficient under normoxia. (A) Gene Ontology biological process terms enriched among predicted 28S-rsRNA-CTC_L20 target genes. (B) Cell cycle distribution of normoxic and hypoxic mPASMCs transfected with control (NC) or 28S-rsRNA-CTC_L20 inhibitor (n=3 independent experiments). (C) Immunoblots of PCNA, CDK1, CDK2, cyclin A and cyclin E (n=6 independent experiments). (D) EdU and Ki67 staining, scored as the percentage of positive cells (n=6 independent cultures per group for quantification). Scale bar, 50 μm. (E) Cell viability under the same conditions (n=6 independent cultures per group). (F) mPASMCs overexpressing 28S-rsRNA-CTC_L20 under normoxia: transcript level, viability, EdU and Ki67 staining, and immunoblots of PCNA, CDK2 and cyclin A (n=6 independent experiments per group). Scale bar, 50 μm. Data are mean±SEM. CDK indicates cyclin-dependent kinase; EdU, 5-ethynyl-2′-deoxyuridine; NC, negative control; PCNA, proliferating cell nuclear antigen.

Conversely, transfection of a 28S-rsRNA-CTC_L20 mimic increased the expression of 28S-rsRNA-CTC_L20 in normoxic mPASMCs (Fig. 2F). Overexpression of 28S-rsRNA-CTC_L20 under normoxia promoted cell viability (Fig. 2F), increased the expression of PCNA, CDK2 and Cyclin A (Fig. 2F, S2C) and increased EdU incorporation and Ki67 positivity (Fig. 2F).

### 28S-rsRNA-CTC_L20 sustains mPASMC proliferation through β-hydroxybutyrate

KEGG pathway enrichment analysis (Metascape) of the predicted target genes of 28S-rsRNA-CTC_L20 showed that the enriched pathways were mainly involved in the cell cycle and lipid metabolism, including fatty acid metabolism (Fig. 3A). We therefore measured the ketone body content in mPASMCs. The BHB content was decreased in hypoxic mPASMCs after silencing 28S-rsRNA-CTC_L20 and increased in normoxic mPASMCs overexpressing 28S-rsRNA-CTC_L20 (Fig. 3B), and the acetoacetate content was decreased in hypoxic mPASMCs after silencing 28S-rsRNA-CTC_L20 (Fig. S3).

**Figure 3.**
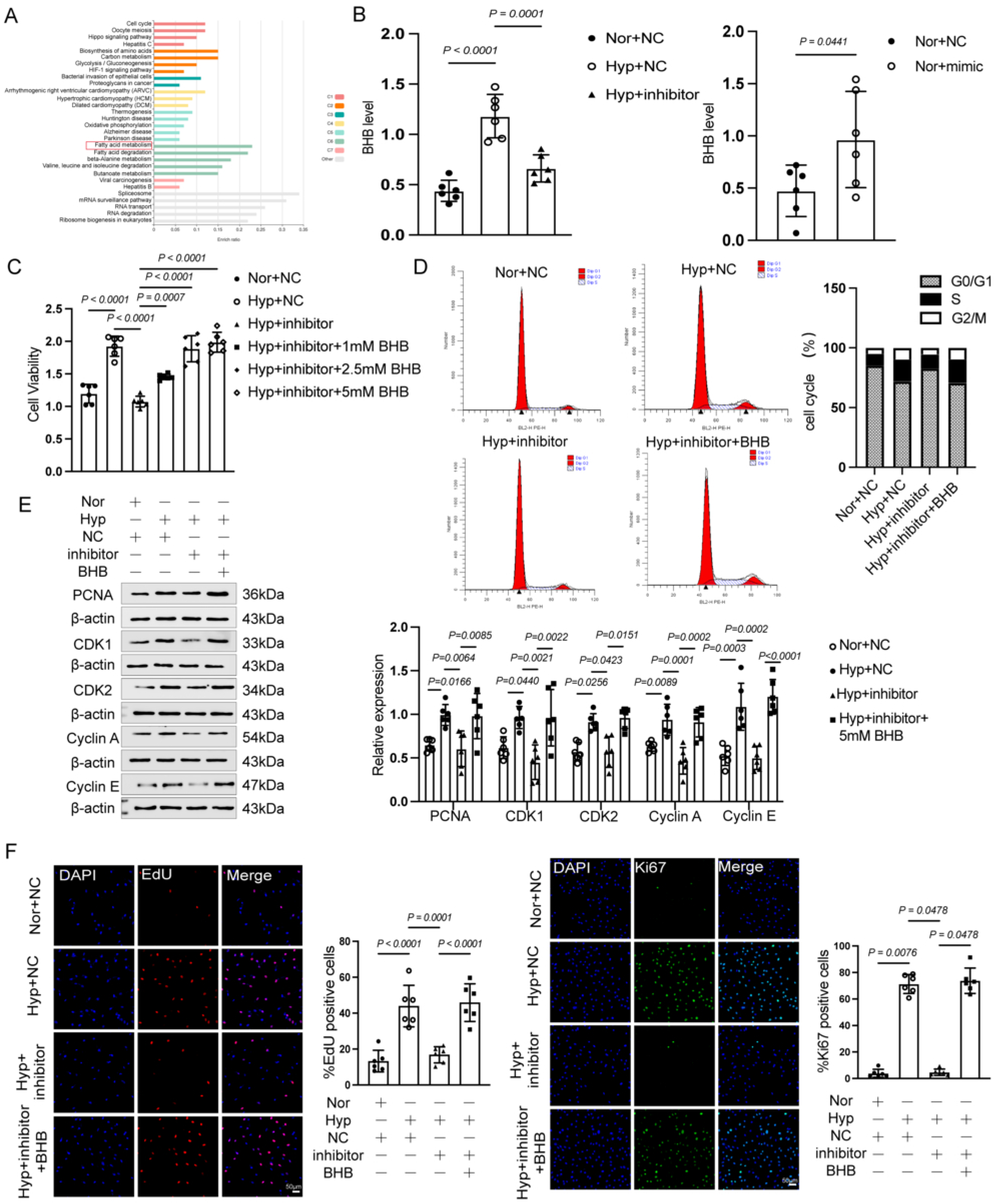
28S-rsRNA-CTC_L20 sustains mPASMC proliferation through β-hydroxybutyrate. (A) KEGG pathways enriched among predicted 28S-rsRNA-CTC_L20 targets (Metascape). (B) BHB content in hypoxic mPASMCs after silencing and in normoxic mPASMCs after overexpression (left and right, respectively; n=6 independent cultures per group). (C) Viability of hypoxic mPASMCs given the inhibitor together with 1, 2.5 or 5 mM BHB (n=6 independent cultures per group). (D) Cell cycle analysis, with representative histograms (n=3 independent experiments). (E) Immunoblots and densitometry for PCNA, CDK1, CDK2, cyclin A and cyclin E (n=6 independent experiments). (F) EdU and Ki67 staining (n=6 independent cultures per group). Scale bar, 50 μm. Data are mean±SEM. BHB indicates β-hydroxybutyrate; KEGG, Kyoto Encyclopedia of Genes and Genomes.

We next examined whether BHB mediated the effect of 28S-rsRNA-CTC_L20 on mPASMC proliferation. The addition of 1 mM, 2.5 mM and 5 mM BHB restored the viability of hypoxic mPASMCs in which 28S-rsRNA-CTC_L20 had been silenced (Fig. 3C). BHB also reversed the inhibition of S-phase entry (Fig. 3D) and restored the expression of PCNA, CDK1, CDK2, Cyclin A and Cyclin E (Fig. 3E) and the proportion of EdU- and Ki67-positive cells (Fig. 3F) in hypoxic mPASMCs treated with the 28S-rsRNA-CTC_L20 inhibitor.

### 28S-rsRNA-CTC_L20 stabilizes the mRNAs of ketone body-synthesizing enzymes through a CTGNNGG motif

We next investigated how 28S-rsRNA-CTC_L20 regulates the generation of BHB (Fig. 4A). Silencing 28S-rsRNA-CTC_L20 reduced the mRNA and protein levels of BDH1, HMGCS2, HMGCL and OXCT1 in hypoxic mPASMCs (Fig. 4B, S4A). Sequence analysis indicated that 28S-rsRNA-CTC_L20 targets the mRNAs of HMGCL, HMGCS2 and BDH1 via its “CTGNNGG” motif (Fig. 4C). In actinomycin D-treated hypoxic mPASMCs, the 28S-rsRNA-CTC_L20 inhibitor accelerated the degradation of BDH1, HMGCS2 and HMGCL mRNA (Fig. 4C), whereas it did not affect the degradation of PLD1 mRNA (Fig. S4B).

**Figure 4.**
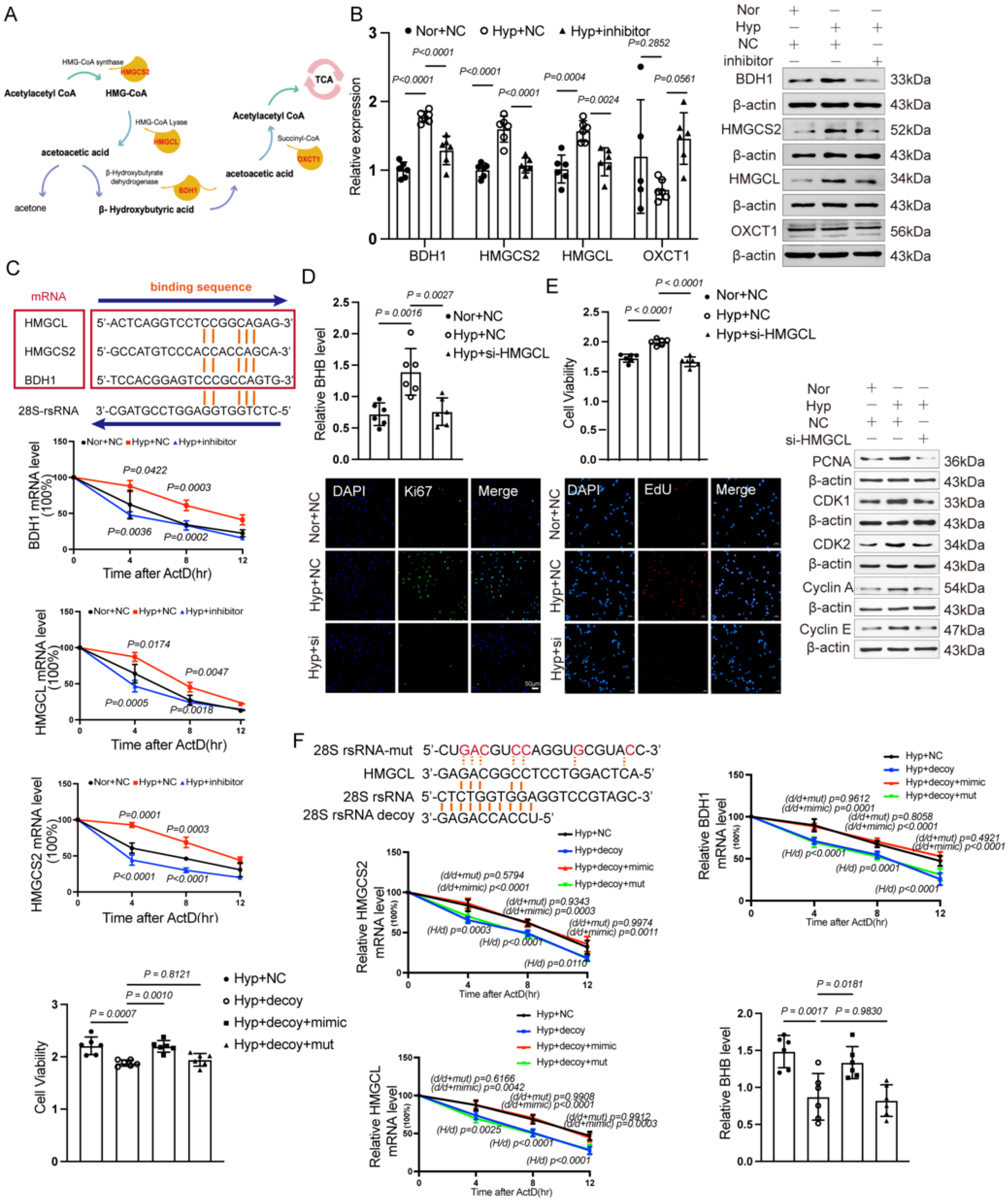
28S-rsRNA-CTC_L20 stabilizes the mRNAs of ketone body-synthesizing enzymes through a CTGNNGG motif. (A) Ketone body synthesis pathway. (B) mRNA and protein levels of BDH1, HMGCS2, HMGCL and OXCT1 in hypoxic mPASMCs after silencing (n=6 independent cultures per group). (C) Predicted base pairing of 28S-rsRNA-CTC_L20 with HMGCL, HMGCS2 and BDH1 mRNAs (top); decay of the three transcripts after actinomycin D (ActD) treatment (bottom; n=6 independent cultures per group). (D) BHB content after HMGCL knockdown (n=6 independent cultures per group). Scale bar, 50 μm. (E) Viability, Ki67 staining, EdU incorporation and immunoblots of cell cycle proteins in the same setting (n=6 independent cultures per group). Scale bar, 50 μm. (F) Mutated 28S-rsRNA-CTC_L20 sequences (top), and decay of HMGCS2, HMGCL and BDH1 mRNAs, viability of hypoxic mPASMCs, together with BHB content in cells cotransfected with decoy and WT or mutant mimic (bottom; n=6 independent cultures per group). Data are mean±SEM. ActD indicates actinomycin D; BHB, β-hydroxybutyrate; WT, wild type.

Knockdown of HMGCL reduced the BHB content in hypoxic mPASMCs (Fig. 4D) and decreased Ki67 positivity, EdU incorporation and cell viability (Fig. 4D, E, S4C), accompanied by reduced expression of PCNA, CDK1, CDK2, Cyclin A and Cyclin E (Fig. 4E, S4D).

To determine whether the effect of 28S-rsRNA-CTC_L20 depends on the predicted binding sequence, we mutated the functional sequence of 28S-rsRNA-CTC_L20 (Fig. 4F). In hypoxic mPASMCs transfected with the 28S-rsRNA-CTC_L20 decoy, the wild-type 28S-rsRNA-CTC_L20 mimic, but not the mutant, slowed the degradation of BDH1, HMGCS2 and HMGCL mRNA and increased the BHB content, Ki67 positivity, EdU incorporation and cell viability (Fig. 4F, S4E). We also examined the endogenous mRNA stability of HMGCL and the relative BHB level in hypoxic mPASMCs transfected with the mRNA decoy of wild-type or mutant HMGCL (Fig. S4F).

### BHB promotes K166 β-hydroxybutyrylation of PKM2 to sustain mPASMC proliferation

To identify downstream effectors of BHB, we analyzed the differentially expressed genes in hypoxic mPASMCs. Several glycolysis-related genes, including Pkm, Slc16a1, Aldoc and Egln1, were upregulated (Fig. 5A), and KEGG pathway enrichment analysis (Metascape) identified glycolysis/gluconeogenesis as the most significantly enriched pathway among the differentially expressed genes (Fig. 5B). Mass spectrometry identified lysine 166 (K166) of PKM2 as a β-hydroxybutyrylation (Kbhb) site (Fig. 5C). Co-immunoprecipitation showed that the total protein level and the Kbhb level of PKM2 were increased in hypoxic mPASMCs and in the lung tissues of patients with PH compared with normoxic mPASMCs and control donors (Fig. 5C, S5A, B).

**Figure 5.**
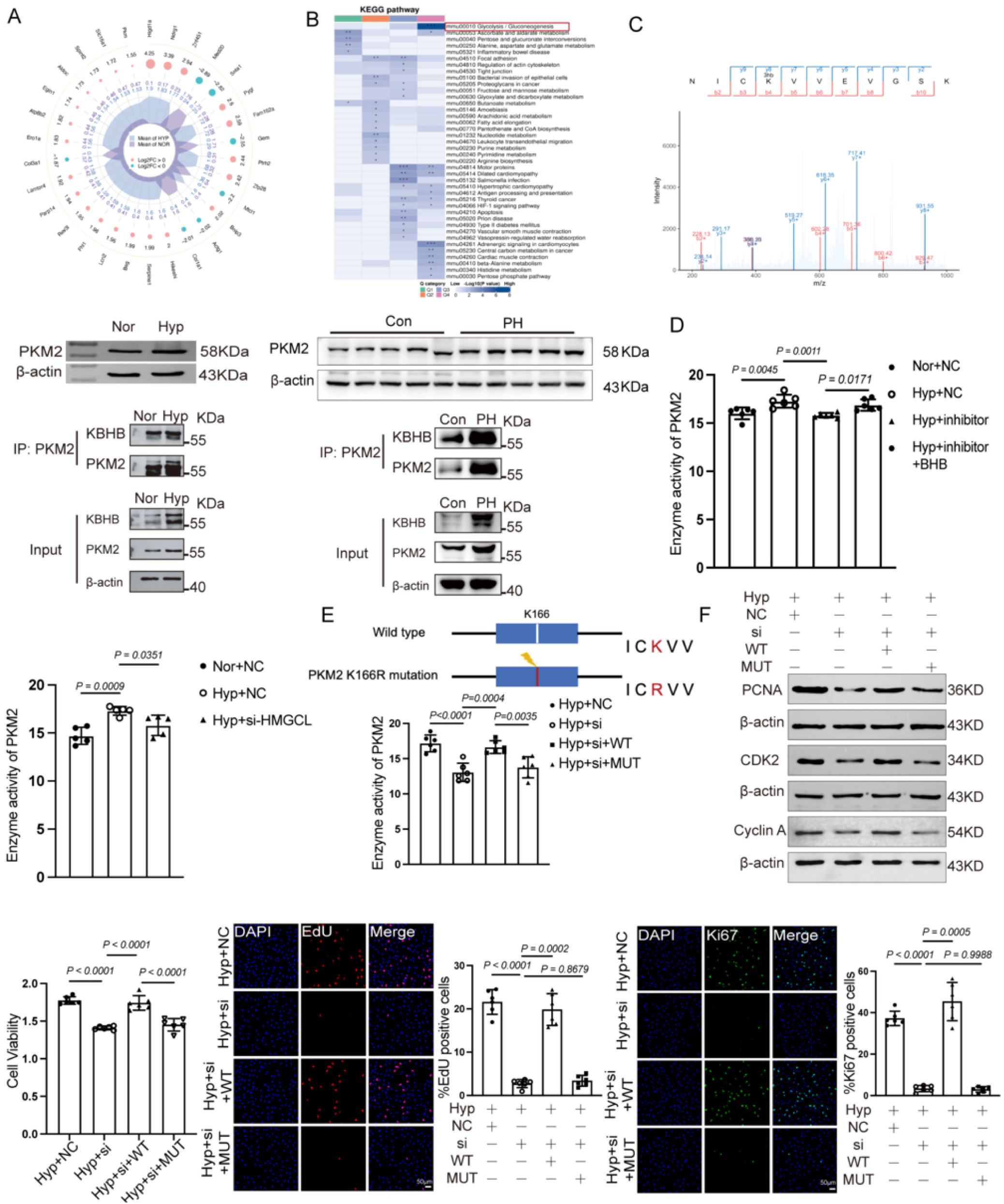
BHB promotes K166 β-hydroxybutyrylation of PKM2 to sustain mPASMC proliferation. (A) Differentially expressed genes in hypoxic versus normoxic mPASMCs, arranged with the enriched Metascape terms. (B) KEGG enrichment of the same gene set. (C) Mass spectra identifying K166 as a Kbhb site in PKM2 (top), PKM2 expression (bottom, n=6 independent experiments; n=5 donors per group for human lung) and PKM2 Kbhb in normoxic and hypoxic mPASMCs and in lung tissue from donors and patients with PH (bottom, n=3 independent experiments; n=3 donors per group for human lung). (D) PKM2 activity in hypoxic mPASMCs after silencing of HMGCL or 28S-rsRNA-CTC_L20, with or without BHB (n=6 independent cultures per group). (E) K166R substitution (top) and PKM2 activity in cells depleted of PKM2 and reconstituted with WT or K166R protein (n=6 independent cultures per group). (F) Immunoblots, viability, EdU incorporation and Ki67 positivity from the same reconstitution experiment (n=6 independent cultures per group). Scale bar, 50 μm. Data are mean±SEM. Kbhb indicates lysine β-hydroxybutyrylation; PKM2, pyruvate kinase M2.

PKM2 enzyme activity was decreased in hypoxic mPASMCs treated with the 28S-rsRNA-CTC_L20 inhibitor or HMGCL siRNA, and was restored by the addition of BHB (Fig. 5D). To determine whether Kbhb of PKM2 at K166 was required, we replaced the lysine at position 166 with arginine (K166R) (Fig. 5E).

In hypoxic mPASMCs transfected with PKM2 siRNA, PKM2 enzyme activity was restored by wild-type PKM2 but not by the K166R mutant (Fig. 5E). Similarly, the expression of PCNA, CDK2 and Cyclin A, cell viability, EdU incorporation and Ki67 positivity were restored by wild-type PKM2 but not by the K166R mutant (Fig. 5F, S5C).

### Knocking down 28S-rsRNA-CTC_L20 attenuates pulmonary hypertension and vascular remodeling in SuHx mice

To evaluate the effect of 28S-rsRNA-CTC_L20 in vivo, SuHx mice were treated with AAV6 carrying a 28S-rsRNA-CTC_L20 antisense oligonucleotide (AAV6-ASO) (Fig. 6A). RT-qPCR confirmed that AAV6-ASO reduced the expression of 28S-rsRNA-CTC_L20 in the lung tissues (Fig. 6A), and FISH combined with immunofluorescence showed reduced expression of 28S-rsRNA-CTC_L20 in the pulmonary arteries, with α-SMA used to localize the smooth muscle layer (Fig. S6A). Knockdown of 28S-rsRNA-CTC_L20 reduced right ventricular systolic pressure (RVSP) (Fig. 6B), prolonged pulmonary artery acceleration time (PAAT) and increased pulmonary artery velocity time integral (PAVTI) (Fig. 6C), and reduced the weight ratio of the right ventricle to the left ventricle plus septum (RV/(LV+Septum)) (Fig. 6D). HE and Masson staining of the lung sections showed that AAV6-ASO attenuated pulmonary vascular remodeling (Fig. 6E), and the protein levels of PCNA, CDK2 and Cyclin A in the lung tissues were reduced (Fig. 6F). HE staining of the heart, liver, spleen and kidney also revealed no obvious pathological changes in the AAV6-ASO-treated mice, and the expression of 28S-rsRNA-CTC_L20 in these tissues was not significantly altered (Fig. S6B, C).

**Figure 6.**
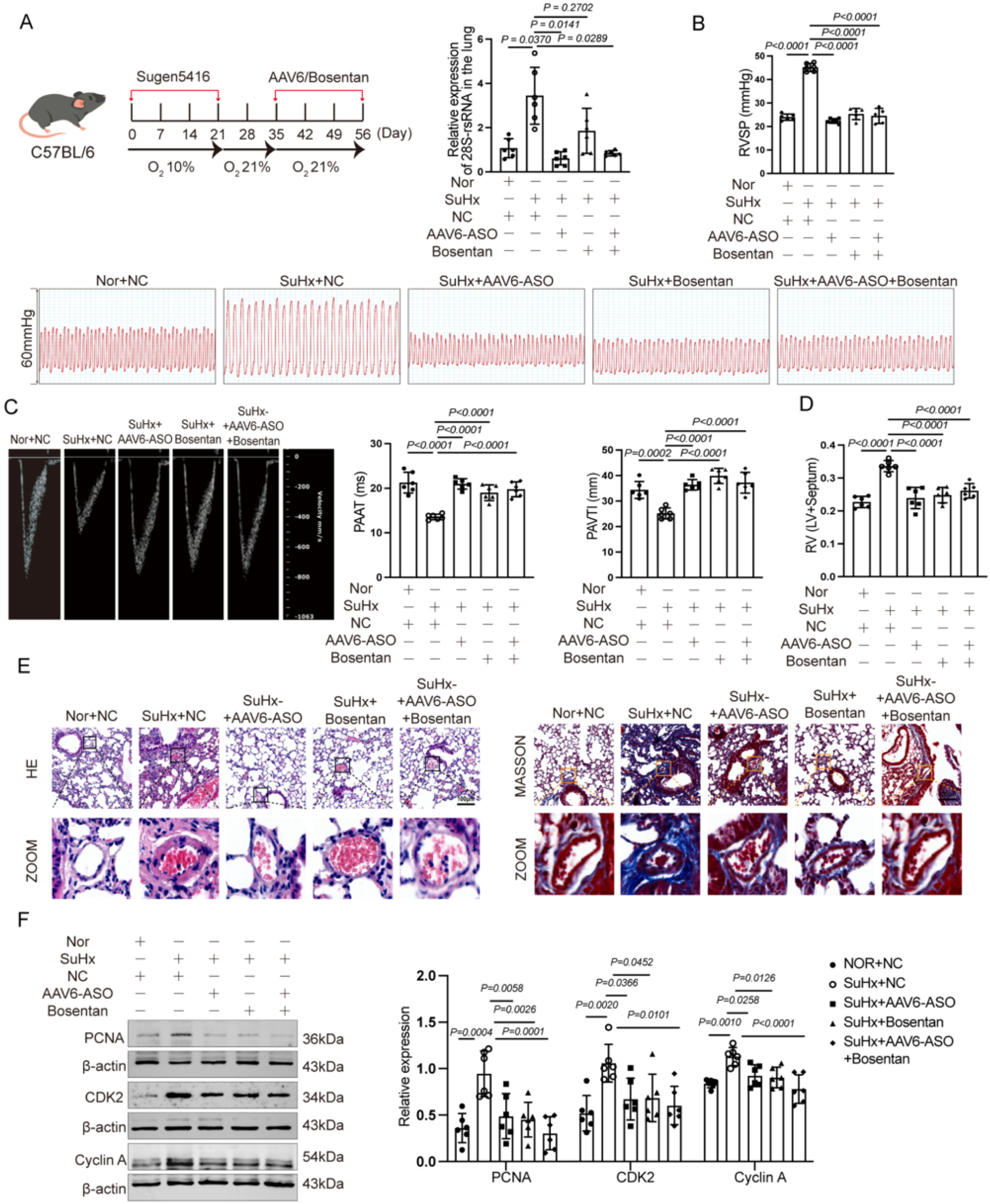
Knocking down 28S-rsRNA-CTC_L20 attenuates pulmonary hypertension and vascular remodeling in SuHx mice. (A) Regimen (SU5416 injection, 3 weeks at 10% O₂, AAV6-ASO or bosentan from day 21, harvest on day 56) and 28S-rsRNA-CTC_L20 in lung tissue (n=6 mice per group). (B) RVSP, with representative pressure tracings (n=6 mice per group). (C) PAAT and PAVTI, with representative echocardiograms (n=6 mice per group). (D) RV/(LV+septum) mass ratio (n=6 mice per group). (E) Hematoxylin-eosin and Masson staining of lung sections, with magnified views (n=3 mice per group). (F) Immunoblots and densitometry of PCNA, CDK2 and cyclin A in lung tissue (n=6 mice per group). Data are mean±SEM. AAV6 indicates adeno-associated virus serotype 6; ASO, antisense oligonucleotide; PAAT, pulmonary artery acceleration time; PAVTI, pulmonary artery velocity time integral; RV/(LV+septum), right ventricle/(left ventricle+septum); RVSP, right ventricular systolic pressure.

## Discussion

Our data identify 28S-rsRNA-CTC_L20 as a hypoxia-inducible, PASMC-enriched small non-coding RNA that couples ketone body production to glycolytic reprogramming and pulmonary vascular remodeling. Several observations support this interpretation. 28S-rsRNA-CTC_L20 was among the most strongly upregulated sncRNAs in PANDORA-seq profiling, was induced by hypoxia in mPASMCs but not in mPAECs, and was increased in the pulmonary arteries, lung tissue and blood of SuHx mice, as well as in the plasma and remodeled pulmonary arteries of patients with PH. Silencing 28S-rsRNA-CTC_L20 suppressed hypoxia-induced mPASMC proliferation, and AAV6-mediated delivery of an antisense oligonucleotide attenuated pulmonary vascular remodeling and hemodynamic impairment in SuHx mice. In addition, 28S-rsRNA-CTC_L20 maintained the stability of BDH1, HMGCS2 and HMGCL mRNA and thereby increased the production of BHB, and BHB supplementation restored the proliferation of cells in which 28S-rsRNA-CTC_L20 had been silenced. Finally, BHB increased β-hydroxybutyrylation of PKM2 and PKM2 enzymatic activity, whereas a PKM2 K166R mutant failed to restore proliferation in PKM2-silenced hypoxic mPASMCs. Together, these data define a 28S-rsRNA-CTC_L20/ketone body/PKM2 β-hydroxybutyrylation axis that promotes PASMC proliferation in PH.

Conventional small RNA sequencing underestimates sncRNAs that carry modifications blocking adapter ligation and reverse transcription. PANDORA-seq overcomes this barrier and has shown that tsRNAs and rsRNAs, rather than miRNAs, predominate in many mammalian samples^18,19^. Functional annotation of rsRNAs has lagged behind that of miRNAs and tsRNAs, and most studies have examined them in the germline, aging and cancer, where individual rsRNAs regulate transcript stability and drug resistance^12^ and circulating rsRNAs have been proposed as diagnostic biomarkers^20^. Our data extend this biology to the pulmonary circulation and provide what we believe is the first evidence that an rsRNA participates in the pathogenesis of PH.

Several classes of sncRNA have been implicated in PH. In pulmonary arterial endothelial cells, tsRNA-3040b promotes glycolysis and proliferation by driving R-loop-dependent transcription of Trim35^21^, and tRF-31-PS5P4PW3FJHP suppresses endothelial proliferation by binding LSM4 and altering EDN1 splicing^22^. Our results agree with these reports in that a hypoxia-regulated sncRNA shapes the metabolic phenotype of pulmonary vascular cells, but they differ in two respects. The first is the cellular compartment: the rsRNA described here is induced in PASMCs and is not changed in PAECs under the same conditions, whereas the tsRNAs studied to date have been characterized mainly in endothelial cells. The second is the direction of the effect on the target transcript. Canonical miRNA and tsRNA mechanisms reduce the stability or translation of their targets, whereas 28S-rsRNA-CTC_L20 slowed the degradation of BDH1, HMGCS2 and HMGCL mRNA in actinomycin D chase experiments. This effect required the predicted complementary sequence, because neither a mutant 28S-rsRNA-CTC_L20 nor a mutated HMGCL binding site reproduced it, and it was not observed for PLD1 mRNA. An mRNA-stabilizing activity of this kind has precedent among tiRNAs that protect their targets from degradation^23^ and contrasts with the miRNA-like repression reported for rsRNA-28S in prostate cancer cells^12^. A single sncRNA class can therefore act through more than one post-transcriptional mechanism, depending on the target and the cellular context.

Increased ketone body availability has been linked to cardiopulmonary disease. Philip et al reported that fatty acid oxidation-derived β-hydroxybutyrate sensitizes TRPV4 and increases intracellular calcium in lung microvascular endothelial cells from SuHx rats, and that a transpulmonary BOHB gradient is detectable in patients with PAH^14^. Our observation that BHB is generated within PASMCs and drives their proliferation is consistent with the idea that BHB in the hypertensive lung is more than an alternative fuel^15^, but the two settings differ. In the endothelium, BHB acts as a signaling metabolite at a calcium channel; in PASMCs it acted intracellularly, and supplying BHB to cells in which 28S-rsRNA-CTC_L20 had been silenced restored viability, cell cycle progression and proliferation-related protein expression. Ketone body metabolism has also been invoked in the adaptation of the right ventricle to pressure overload^24^, and pharmacological enhancement of ketone body metabolism improved right heart function in experimental PAH^25^. Whether the ketogenic program described here in PASMCs also operates in the right ventricle, and whether the two are linked through circulating ketone bodies, remains to be determined.

Aerobic glycolysis is a hallmark of remodeled pulmonary vessels and of the proliferative, synthetic PASMC phenotype^26,27^. Earlier work has placed PKM2 at the center of this reprogramming, largely at the level of RNA processing: Caruso et al identified miR-124 as a regulator of endothelial glycolysis acting through PTBP1 and PKM2^28^, and Lu et al reported that reduced CircSMOC1 promotes metabolic reprogramming via PTBP1 and miR-329-3p in PAH rats^5^. Our results add a post-translational layer to this scheme. β-hydroxybutyrylation of K166 increased PKM2 enzymatic activity, and replacing that lysine with arginine abolished the rescue of proliferation in PKM2-silenced hypoxic mPASMCs even though the protein was expressed. Guo et al. reported that reactive oxygen species inhibit PKM2 in PAH^29^, which is the opposite of the Kbhb-dependent activation seen here. PKM2 activity in the pulmonary vasculature therefore appears to be controlled by opposing signals: an oxidative one that suppresses it and a ketone body-dependent modification that enhances it. The net glycolytic and proliferative output reflects the balance between the two. Additional glycolytic checkpoints in PASMCs, including mitochondrial pyruvate import and lactate generation, have been described^30,31^, and the contribution of the Kbhb-dependent route to overall glycolytic flux remains to be quantified.

Lysine β-hydroxybutyrylation was first described as a histone mark induced by ketogenesis^32^ and is now recognized as a modification of non-histone proteins through which ketone body availability is translated into changes in protein function^33,34^. K166 of PKM2 has not previously been reported as a Kbhb site. Our data indicate that this modification is not inert: it was increased in hypoxic mPASMCs and in the lungs of patients with PH, and mutating the site abolished the rescue of both PKM2 activity and proliferation. Kbhb therefore provides a mechanistic link between a lipid-derived metabolite and the glycolytic enzyme that sustains the proliferative phenotype of PASMCs.

Two aspects of our findings are relevant to biomarker and therapeutic development. 28S-rsRNA-CTC_L20 was detectable in plasma and was increased in patients with PH, consistent with the growing interest in circulating sncRNAs as minimally invasive biomarkers in PAH^35^ and with reports of plasma rsRNA signatures in cancer^20^. Knockdown with AAV6-ASO also reduced pulmonary vascular remodeling without pathological changes in the heart, liver, spleen or kidney, which adds to the evidence that oligonucleotide-based interventions are feasible in PH^36^.

Several limitations should be considered. The SuHx model does not reproduce the full spectrum of human PAH pathology, including plexiform lesions. Human lung tissue and plasma were obtained from a small number of subjects, and the human data are therefore preliminary. We did not identify the nuclease or processing pathway that generates 28S-rsRNA-CTC_L20 under hypoxia, nor did we determine whether the shift of the transcript toward the cytoplasm is required for its stabilizing activity. Only one Kbhb site was interrogated, and other Kbhb substrates may contribute to the proliferative phenotype. The BHB concentrations required to restore proliferation were in the millimolar range, and the relative contribution of the ketogenic route to intracellular BHB in vivo remains to be established. We also did not examine the roles of other post-translational modifications of PKM2 in proliferation. Finally, we cannot exclude additional mRNA targets of 28S-rsRNA-CTC_L20 beyond BDH1, HMGCS2 and HMGCL. Future work should define the biogenesis of this rsRNA and determine whether circulating 28S-rsRNA-CTC_L20 tracks disease severity or treatment response. Whether inhibiting the ketogenic program or PKM2 Kbhb attenuates established disease also needs to be tested.

## Perspectives

Excessive proliferation of pulmonary artery smooth muscle cells remains the principal driver of vascular remodeling in PAH, and current vasodilator therapies do not reverse it. This study shows that rRNA-derived small RNAs, a class of small non-coding RNA largely overlooked in cardiovascular research, can set the metabolic tone of the pulmonary artery wall. 28S-rsRNA-CTC_L20 stabilizes the mRNAs of ketone body-synthesizing enzymes, raises intracellular β-hydroxybutyrate and licenses the glycolytic enzyme PKM2 through β-hydroxybutyrylation at K166. Two implications follow. Ketone bodies in the hypertensive lung act as signaling molecules that modify protein function rather than merely as fuels, which may explain why interventions that alter ketone body handling have cardiovascular effects beyond energetics. The rsRNA-ketogenesis-Kbhb axis also offers several points at which vascular remodeling might be interrupted, including the rsRNA itself, HMGCL-dependent ketone body production, and the Kbhb modification. Future work should establish how hypoxia generates this rsRNA and whether plasma 28S-rsRNA-CTC_L20 or its associated ketone body signature stratifies patients. Whether the axis is active in other forms of PH and in the pressure-loaded right ventricle also remains open.

## Supplementary Material

Expanded Methods, Figures S1-S6

## Author Contributions

H.Y. and D.Z. designed research; A.L., Y.P., C.L., J.Z., X.L., Y.W., H.Z., Z.B., X.Z., Y.X. and Y.J. performed research; H.Y. and D.Z. contributed new reagents/analytic tools and funding support; A.L., Y.P., C.L., J.B., H.S. and Y.W. analyzed data; and H.Y., A.L. and H.T. wrote the paper. **Competing Interests.** The authors declare that they have no competing interests. **Data and materials availability:** All data needed to evaluate the conclusions in the paper are present in the paper and/or the Supplementary Materials. The mechanistic diagram was created with Figdraw.

## Novelty and Relevance What Is New?

PANDORA-seq identified the rRNA-derived small RNA 28S-rsRNA-CTC_L20 as a hypoxia-induced sncRNA in pulmonary artery smooth muscle cells; it is increased in SuHx mouse lungs and blood and in the plasma and pulmonary arteries of patients with pulmonary hypertension.

28S-rsRNA-CTC_L20 binds and stabilizes the mRNAs of the ketone body-synthesizing enzymes BDH1, HMGCS2 and HMGCL, increasing β-hydroxybutyrate production.

β-hydroxybutyrate drives proliferation through β-hydroxybutyrylation of PKM2 at lysine 166, and an AAV6-delivered antisense oligonucleotide against the rsRNA attenuates pulmonary hypertension in SuHx mice.

## What Is Relevant?

Pulmonary hypertension is characterized by metabolic reprogramming and excessive proliferation of pulmonary artery smooth muscle cells; this study links a previously unstudied small RNA class to that phenotype.

Ketone bodies and lysine β-hydroxybutyrylation are established metabolic signals, but neither has previously been placed upstream of vascular remodeling in the pulmonary circulation.

## Clinical / Pathophysiological Implications?

Circulating 28S-rsRNA-CTC_L20 is detectable and elevated in patients with pulmonary hypertension, supporting further evaluation of rRNA-derived small RNAs as minimally invasive biomarkers.

Targeting the rsRNA, HMGCL-dependent ketogenesis or PKM2 β-hydroxybutyrylation may offer routes to anti-remodeling therapy that complement existing vasodilator treatment.

